# Notch-mediated thyroid hormone regulation of skin development in the zebrafish caudal fin

**DOI:** 10.64898/2026.03.20.713269

**Authors:** Toshiaki Uemoto, Melody Autumn, Sarah K McMenamin

## Abstract

Thyroid hormone (TH) is a systemic regulator of vertebrate development, yet its role in the maturation of the stratified skin remains poorly defined. Using the skin at the edge of the zebrafish caudal fin, we defined the trajectory of epidermal maturation during the transition from juvenile to adult. We found the peripheral edge (PE) of the fin exhibits positive allometric expansion that is dependent on TH: in thyroid-ablated, hypothyroid backgrounds, the growth of the PE is limited. We showed that TH drives normal PE growth by stimulating both cell proliferation and hypertrophy. Further, we demonstrated that TH acts upstream of the Notch pathway to regulate growth of the PE. While TH signaling machinery is broadly expressed throughout the fin, Notch pathway activation is localized and highly enriched in the PE. Repressing Notch activity prevented PE expansion, while upregulating Notch in a hypothyroid background was sufficient to increase hypertrophy and partially rescue PE expansion. By identifying Notch as a region-specific effector of TH-driven hypertrophy, our findings show a mechanism by which systemic endocrine signals are translated into local tissue morphogenesis.

**Graphical abstract:** 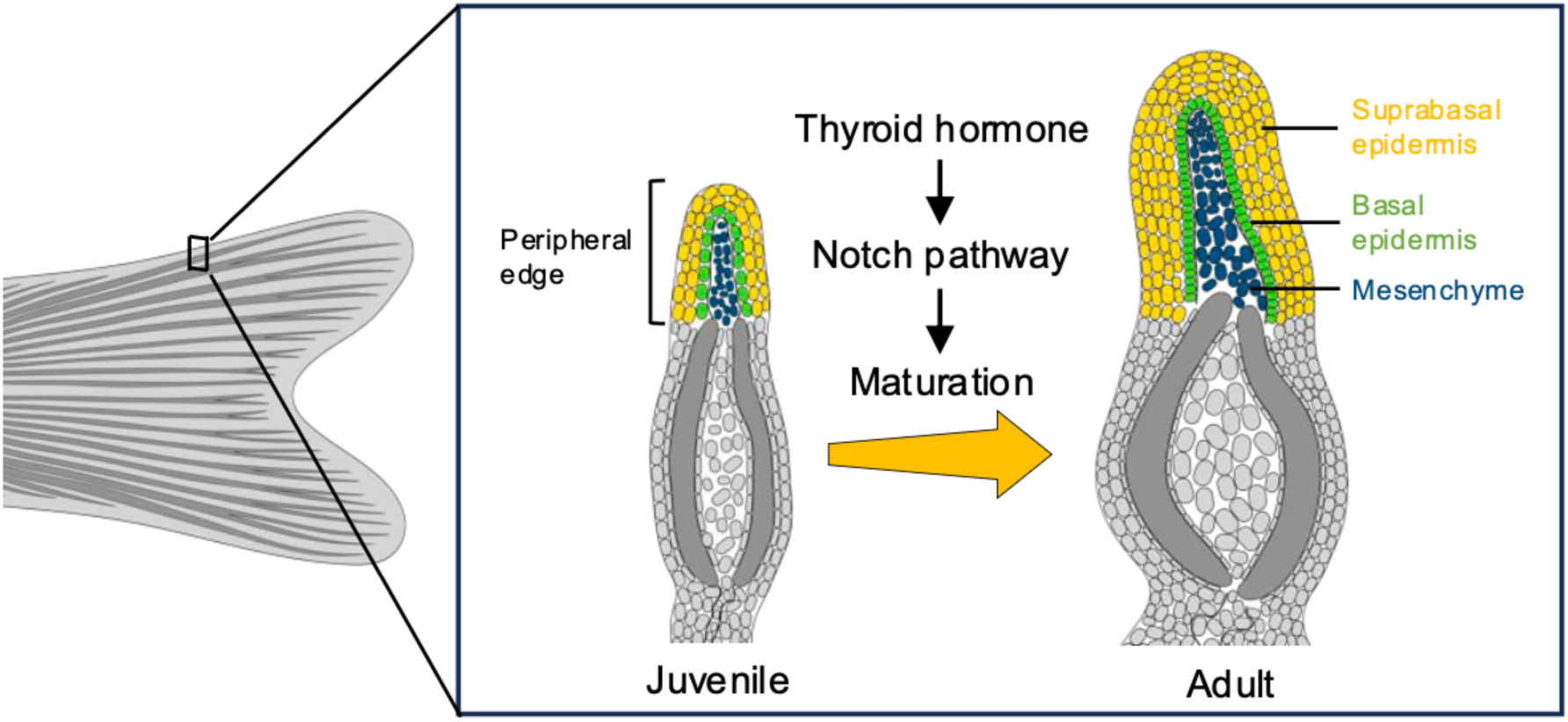

## Introduction

Thyroid hormone (TH) is a crucial regulator of vertebrate post-embryonic development, which coordinates diverse biological transitions including metamorphosis in fish and amphibians, and the maturation of multiple organ systems in mammals^1–5^. One understudied target of TH-stimulated morphogenesis is the vertebrate integument—the organ system comprising the skin and skin appendages. During the post-embryonic stages of teleost and amphibian development, TH drives dramatic structural maturation and thickening of the epidermis, the outermost layer of the skin^6–8^. In mammals, TH is required to maintain epidermal growth and appropriate thickness during post-embryonic stages^9,10^.

Tissues can grow by increasing the number of cells (cell proliferation leading to hyperplasia) and growth of the cells themselves (cell expansion leading to hypertrophy)^11^. In the mammalian epidermis, TH stimulates cell proliferation and maintains epidermal thickness by regulating proliferation and ultimate cell number^9,12,13^. While TH is known to drive cellular hypertrophy in certain tissues, including cartilage^14,15^ and the scale-forming cells of the teleost integument^16^, its potential role in the hypertrophy of the epidermis itself remains unexplored. Like mammals, zebrafish have a stratified skin structure composed of layered epidermis and mesenchyme, and zebrafish are considered a promising model for studying skin development and homeostasis^17–20^. In the zebrafish flank, skin development is intimately coupled with the highly organized formation of scales^21^, a TH-driven process^16,22^. To investigate the fundamental processes underlying TH-mediated epidermal maturation without the anatomical complexity of the scales, we focused on the non-skeletal regions of the caudal fin, which maintain a stratified architecture but which never undergo squamation^23,24^.

The zebrafish caudal fin is composed of segmented skeletal fin rays, surrounded by skin tissue, specifically the inter-ray (IR) tissue, and the non-skeletal peripheral edge (PE) at the upper and lower margins of the fin^25^ (Fig. 1A). The skin of the fins is composed of stratified epidermal layers (suprabasal and basal) and underlying mesenchyme. The PE and IR regions are beside or between the bony fin rays, and their accessibility and simple, relevant structure makes them an attractive model for studying skin maturation.

**Figure 1.**
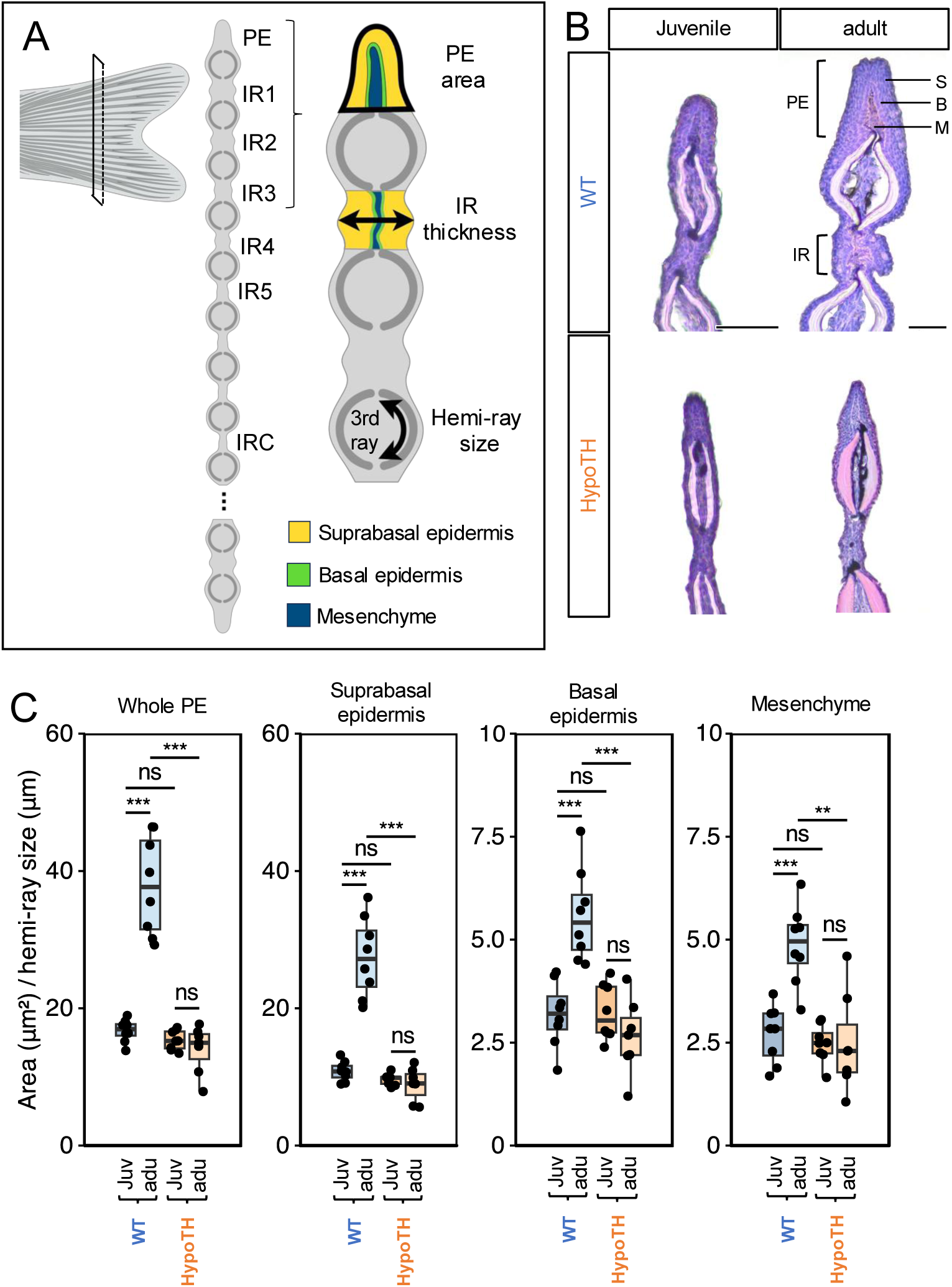
TH is required for the stage-specific allometric growth of the peripheral edge. (A) Schematic illustration of the adult zebrafish caudal fin indicating the plane of cross-section (dashed line). The cross-sectional diagram defines the anatomical regions: peripheral edge (PE), inter-ray (IR) tissues (IR1-5 and IRC), and fin rays. The magnified views of the PE and IR show the tissue organization and color-coding used for analysis. (B) Representative Hematoxylin and Eosin (HE) stained cross-sections of the peripheral rays in wild-type (WT) and TH-deficient (HypoTH) fish at juvenile and adult stages. Scale bars: 0.05 mm. (C) Quantification of the cross-sectional area of the PE, normalized to the hemi-ray width. (D) Quantification of the normalized areas of whole PE and specific tissue layers within the PE. Data in (C) and (D) are presented as box plots showing the median, quartiles, and individual data points. Asterisks indicate significant differences between stages within the stage or condition (**P < 0.01, ***P < 0.001; ns, not significant).

In this study, we demonstrated that TH is essential for the allometric expansion of the fin PE tissues, with the hormone stimulating both cell proliferation and hypertrophy. We found that TH signaling regulates the structural maturation of the fin epidermis during post-embryonic development in a region-specific manner via the Notch pathway. By identifying Notch as a region-specific effector of TH-driven hypertrophy, our findings provide insight into how systemic endocrine signals may be translated into local tissue morphogenesis.

## Results

### Maturation of the peripheral edge tissue of the caudal fin

To establish a baseline for normal post-embryonic maturation of the fin soft tissues, we asked how these regions changed during the transition from juvenile to adult in wild-type (WT) zebrafish. We histologically evaluated transverse sections of fins from both juvenile and adult using hematoxylin and eosin staining (Fig. 1B). At the juvenile stage, zebrafish already possess the definitive forked shape characteristic of adults^26,27^ and the fin epidermis exhibits a stratified structure (Fig. 1B, top row). To detect changes in tissue composition during the transition from juvenile to adult, we measured the cross-sectional area of the PE at each stage (Fig. 1A). We found that while the fundamental tissue architecture was already established by the juvenile stage (Fig. 1B, top row), the normalized PE area increased considerably as fish entered adulthood (Fig. 1B and C). The PE exhibits positive allometric growth during maturation, growing relatively faster than the bony rays widen. To measure the tissue-level contributions to this growth, we analyzed the constituent layers of the PE (suprabasal epidermis, basal epidermis, and mesenchyme) using two metrics: the normalized absolute area of each layer and the fraction of the area. While the normalized areas of all tissue layers significantly increased from juvenile to adult stages (Fig. 1C), their relative proportions were maintained (Fig. S1A and B). This indicates that the developmental expansion of the PE is a uniform scaling of all layers.

### TH signaling machinery is broadly expressed in the caudal fin

TH is synthesized throughout larval and juvenile development^28–30^ and is known to affect fin morphogenesis^31–34^. The canonical TH signaling pathway involves cellular uptake by transporters (e.g., Mct8, Mct10), intracellular conversion of T4 to the more genomically active form T3 by deiodinases (Dios), and subsequent transcriptional regulation as T3 binds to nuclear TH receptors (TRs)^35–37^. We assessed whether TH signaling is active in fin tissues by re-analyzing expression data from intact adult caudal fin tissues^33^. This analysis confirmed that key pathway components, including *mct10*, *dio1*, *dio2, thraa* and *thrb*, are each expressed in the fin tissue (TPM ≥ 1 in all three biological replicates; Fig.S2). Immunohistochemical analysis of select TH pathway components detected Mct10, Dio1, Dio2 and Thra broadly in epidermal cells throughout the fin (Fig. S3). Notably, Mct10 and Dio1 were present in the mesenchyme of fin rays (Fig. S3). Collectively, these findings suggest that the fin soft tissues are capable of actively responding to TH signaling.

### TH is required for allometric expansion of the peripheral edge tissue

To determine whether TH is required for adult PE growth, we evaluated these tissues in thyroid-ablated backgrounds lacking TH (HypoTH)^5,38^. Fins from both juvenile and adult HypoTH fish exhibited the stratified organization of epithelial and mesenchymal cells (Fig. 1B, bottom row). Furthermore, the normalized PE area in juvenile HypoTH fish was indistinguishable from that of WT (Fig. 1C). However, HypoTH fish failed to increase the PE area during the transition from juvenile to adult stages, resulting in a significantly smaller adult PE size compared to WT (Fig. 1C). These results suggest that TH is required for the stage-specific expansion of the PE. Layer-specific analysis revealed that while the normalized areas of all layers were smaller relative to those in adult WT fins (Fig. 1C, Whole PE). Fins of HypoTH adults showed no significant changes in the area fractions of the constituent layers compared to WT adults (Fig. S1A and B). Together, these findings indicate that while TH is not required for basic tissue stratification or the maintenance of relative tissue proportions, it is required for PE expansion.

### TH regulates PE growth through both proliferation and hypertrophy

To determine whether changes in cell number and/or cell size contribute to the TH-mediated growth of the PE, we assessed total cell number by counting DAPI-stained nuclei within each layer. Following a previously described method^39^, we calculated the average cell size from the number of nuclei per unit of area to determine the physical space occupied by each cell; a decrease in cell density was considered to indicate hypertrophy. We compared cell density in fins from WT and HypoTH backgrounds. In juveniles, we observed no significant differences in cell number or density between the two groups (Fig. 2A-C). However, at the adult stage, fins from HypoTH backgrounds showed reduced cell counts across all layers, with a higher cell density in the suprabasal epidermis compared to fins from WT backgrounds (Fig. 2A-C). We tested if this reduction in cell number resulted from impaired proliferation, identifying mitotic cells using an antibody to phosphohistone H3 (PH3). In fins from both WT and HypoTH backgrounds, mitotic cells were restricted to the suprabasal epidermis. There were significantly fewer mitotic cells in fins of HypoTH juveniles compared to WT juveniles; however, no significant difference was detected when adult stages were compared (Fig. 2D). Collectively, these results suggest that TH increases both cell number and cell size to promote PE growth during the transition from juvenile to adult.

**Figure 2.**
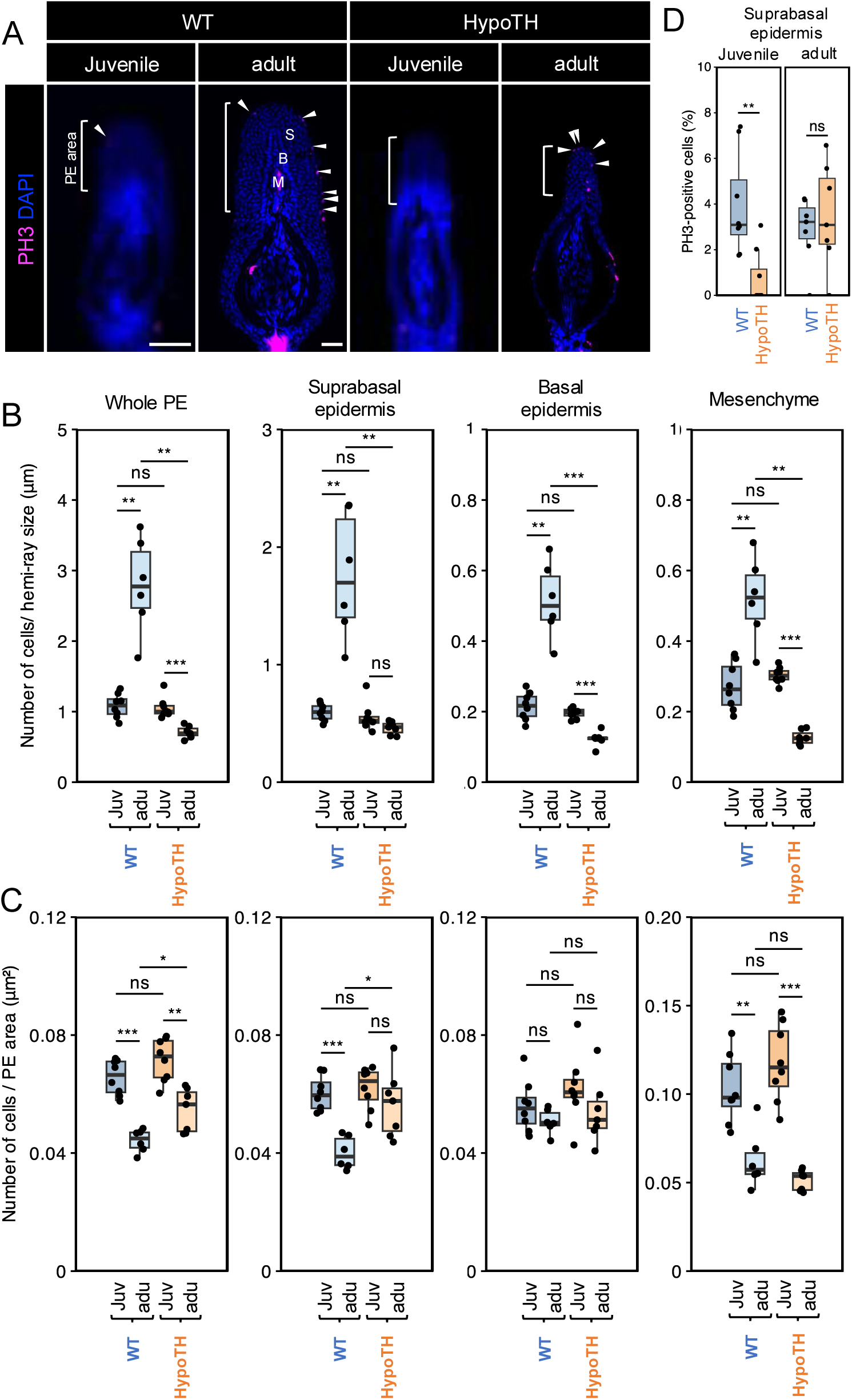
Thyroid hormone promotes PE growth through cell proliferation and hypertrophy. (A) Immunohistochemical detection of PH3 in cross-sections of the peripheral edge (PE) in WT and HypoTH fins. Nuclei are counterstained with DAPI. White arrowheads indicate PH3-positive cells. Scale bars: 20 µm. (B) Total cell number normalized to the hemi-ray size in WT and HypoTH fins. (C) Cell density (cell number per cross-sectional area) in WT and HypoTH fins. (D) Proportion of PH3-positive cells in the suprabasal epidermis. Data are presented as box plots. Asterisks indicate significant differences (**P < 0.01, ***P < 0.001; ns, not significant). Abbreviations: S, suprabasal epidermis; B, basal epidermis; M, mesenchyme.

### TH-dependent maturation of the inter-ray epidermis

Given that key TH pathway components were detectable in the IR (Fig. S3), we asked if TH was required to regulate the IR tissues between the bony rays, or if these effects were limited to the PE tissue at the edge of the fin. In WT fish, IR thickness increased proportionally with fin ray width as fish developed (Fig. S1C top row and D). In contrast, IR tissues in HypoTH contexts failed to maintain this proportionality, exhibiting a relative reduction in IR thickness between the juvenile and adult stages; this resulted in a significantly thinner adult IR compared to that in WT (Fig. S1C bottom row and D). These results indicateTH also regulates the maturation of the IR tissues in addition to the PE.

### TH signaling acts upstream of the Notch pathway in the fin epidermis

Notch signaling has an established role in skin development, regulating epidermal remodeling in amphibians^40^ and controlling keratinocyte proliferation and differentiation in mammals^41^. While Notch signaling regulates blastema cell proliferation and differentiation during zebrafish fin regeneration^42,43^, its function in the structural maturation and homeostatic maintenance of the adult fin epidermis remains unknown. Our preliminary transcriptomic observations^33^ showed a downregulation of Notch pathway components in HypoTH fins (Fig. S4A). Based on these data, we hypothesized that TH acts as an upstream regulator of Notch signaling to control fin epidermal expansion. To visualize Notch signaling *in vivo*, we utilized a Notch activity reporter line *Tg(Tp1bglob:eGFP)*^44^, which drives GFP expression under the control of a Notch-responsive element. Under WT conditions, robust Notch activity was detectable in the PE region (Fig. 3A-A’). Transverse sections revealed Notch signal localization to the suprabasal cells adjacent to the basal epidermis (Fig. 3A’). Notch activity was markedly decreased in HypoTH fins (Fig. 3B and S4C). To test if the attenuation of Notch signaling was a direct consequence of TH deficiency, we treated adult HypoTH fish with exogenous TH (in the form of T4) for 6 days. This supplementation partially restored Notch activity in the periphery of the fin (Fig. 3C and S4C), consistent with direct regulation.

**Figure 3.**
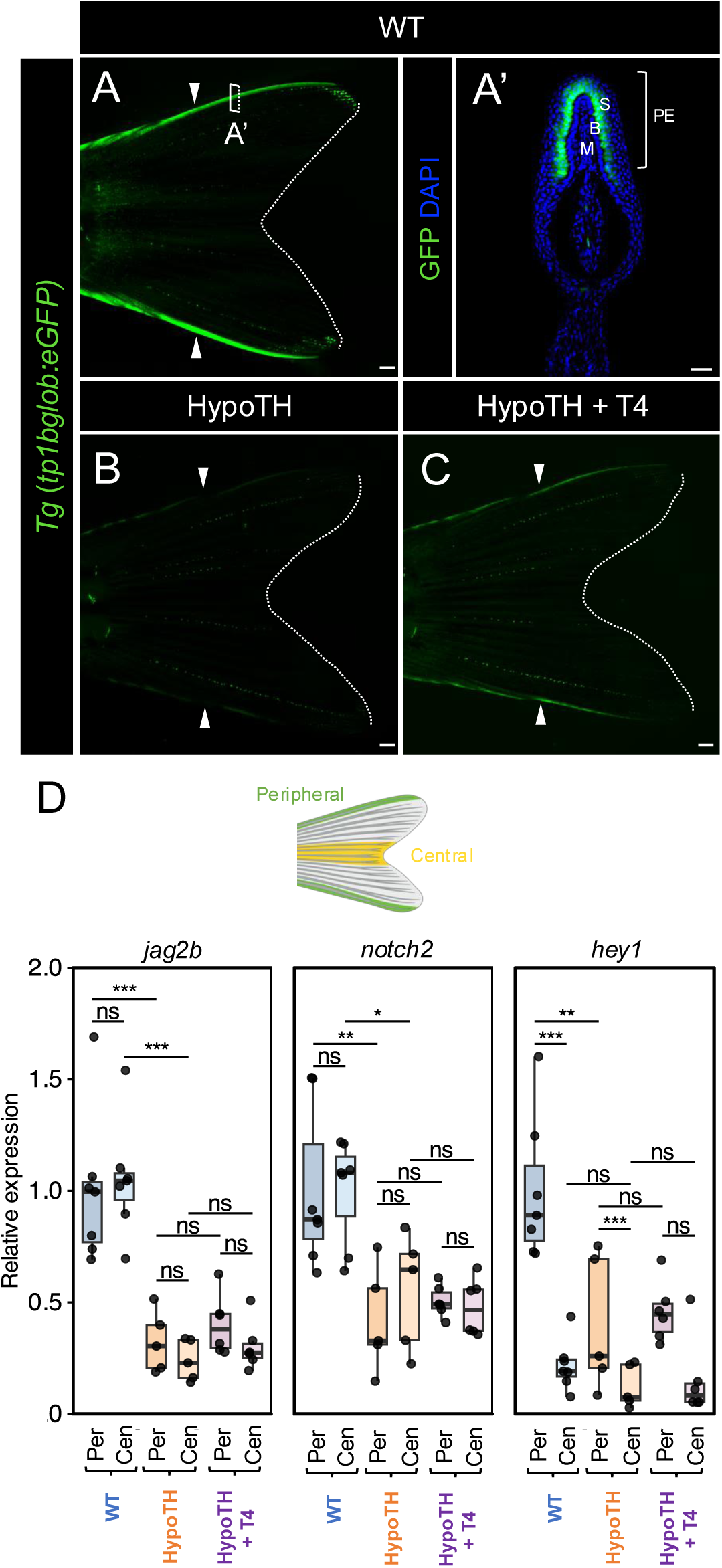
TH-dependent Notch activity in the peripheral fin tissue. (A–C) Live imaging of the caudal fin using the *Tg(Tp1bglob:eGFP)* Notch reporter line. Comparisons are shown for WT (A), HypoTH (B), and T4-supplemented HypoTH (C) conditions. Arrowheads indicate regions of Notch activity at the peripheral margin. All images were acquired using the same imaging settings and exposure times. (A’) Immunohistochemical detection of GFP on the transverse section of a *Tg(Tp1bglob:eGFP)* fin. Notch activity is localized to the suprabasal cells in the PE region. (D) qPCR analysis of *jag2b*, *notch2*, and *hey1* expression levels in peripheral (Per) and central (Cen) sampled regions, as illustrated in the scheme. Data are presented as box plots. Asterisks indicate significant differences (**P < 0.01, ***P < 0.001; ns, not significant). Scale bars: (A–C) 1 μm, (A’) 20μm. Abbreviations: S, suprabasal epidermis; B, basal epidermis; M, mesenchyme; PE, peripheral edge.

To identify specific Notch pathway components targeted by TH signaling, we re-examined our published transcriptome^33^. Genes expressed above the threshold (TPM ≥ 1) included *hey1*, *notch2*, and *jag2b*, which were all significantly downregulated in fin tissues from HypoTH fish compared to WT fish (Fig. S4B). To investigate these targets in a region-specific manner, we quantified their expression by quantitative reverse transcriptase PCR (qRT-PCR) in different dissected regions of fin tissues from adult fish (see cartoon, Fig 3D). In WT fins, *jag2b* and *notch2* exhibited no significant difference in expression between the peripheral and central tissues; however, *hey1* expression was significantly higher in the peripheral tissues compared to the central tissues of WT (Fig. 3D), mirroring the regional activity of the Notch reporter transgenic line (Fig 3A). We further corroborated this spatial distribution using Hybridization Chain Reaction (HCR), which confirmed that *hey1* transcripts were specifically enriched in the PE region and all fin rays in a WT background (Fig. S5A-A’).

To test the TH-dependence of gene expression, we compared transcript abundance in backgrounds that were HypoTH and HypoTH rescued with T4. We confirmed that the expression of these components is TH-dependent: in HypoTH fish, *jag2b* and *notch2* levels declined significantly across both peripheral and central regions, while the peripheral expression of *hey1* was also significantly downregulated (Fig. 3D). However, T4 treatment did not significantly restore expression (Fig. 3D). Consistently, HCR in situ targeting *hey1* showed no detectable recovery of its peripheral expression following T4 supplementation (Fig. 3D and S5B, C). Nonetheless, the depressed expression in HypoTH fins suggests that TH indeed acts as an upstream regulator of Notch across the fin, and specifically maintaining high Notch activity within the peripheral regions.

### Position at the periphery of the fin is not sufficient to induce Notch activity

While TH signaling components are produced throughout the fin tissues (Fig. S3), robust Notch activity was predominantly observed in the PE (Fig. 3A). We reasoned that this spatial restriction might result from location-specific mechanical stimuli, as the fin margins experience significant hydrodynamic stress during swimming^23^. To test whether the mechanical stress of being at the edge of the fin is sufficient to activate Notch signaling, we amputated the Notch-active peripheral-most rays in *Tg(Tp1bglob:eGFP)* fish, thereby exposing the internal ray (2nd most-peripheral ray) to the marginal hydrodynamic environment (Fig. S6A). As the peripheral ray regenerated, the ray was repeatedly trimmed every several days. Through two weeks of continuous exposure, the 2nd peripheral ray did not exhibit detectable increases in Notch activity (Fig. S6B-E). These results suggest that hydrodynamic stress at the periphery of the fin is not sufficient to induce Notch activation.

### Notch inhibition prevents PE expansion

Given that TH regulates PE size and modulates Notch activity, we hypothesized that TH controls PE growth via Notch signaling. We first tested this by inhibiting Notch in a WT background. To test if Notch inhibition alone is sufficient to recapitulate the HypoTH phenotype, we suppressed Notch signaling using a heat-shock inducible dominant-negative Mastermind-like (dnMAML) line^45^. To ensure that this approach directly affected Notch signaling, we confirmed that dnMAML activation indeed decreased expression of the Notch target *hey1* in the peripheral rays (Fig. S7). To evaluate how sustained Notch inhibition affects PE growth, we performed daily heat shock treatments from 10 days post-fertilization (dpf; approximately the onset of caudal fin ray ossification) through adulthood. Histological assessment confirmed that while fins expressing dnMAML retained the stratified architecture of the epidermis and mesenchyme (Fig. 4A), the normalized PE area was significantly reduced (Fig. 4B). Layer-specific analysis revealed that this PE size reduction resulted from a uniform decrease in the size of all layers relative to WT (Fig. 4B). Concomitantly, the area fractions of these layers showed no significant difference between WT and fish expressing dnMAML (Fig. S8A and B). To determine whether reduced PE size in dnMAML fish resulted from a decrease in cell number or cell size, we quantified DAPI-stained nuclei (Fig. S8C). Total cell numbers did not differ significantly between WT and dnMAML fins (Fig. 4C). In contrast, fins expressing dnMAML exhibited a significantly higher cell density specifically in the suprabasal layer compared to WT fins (Fig. S8C and 4D). Given that adult HypoTH fins showed reduced IR thickness compared to WT (Fig. 1B and C), we also analyzed IR thickness in the fish expressing dnMAML. The thickness of the IR region remained unaffected (Fig. S8D and E).

**Figure 4.**
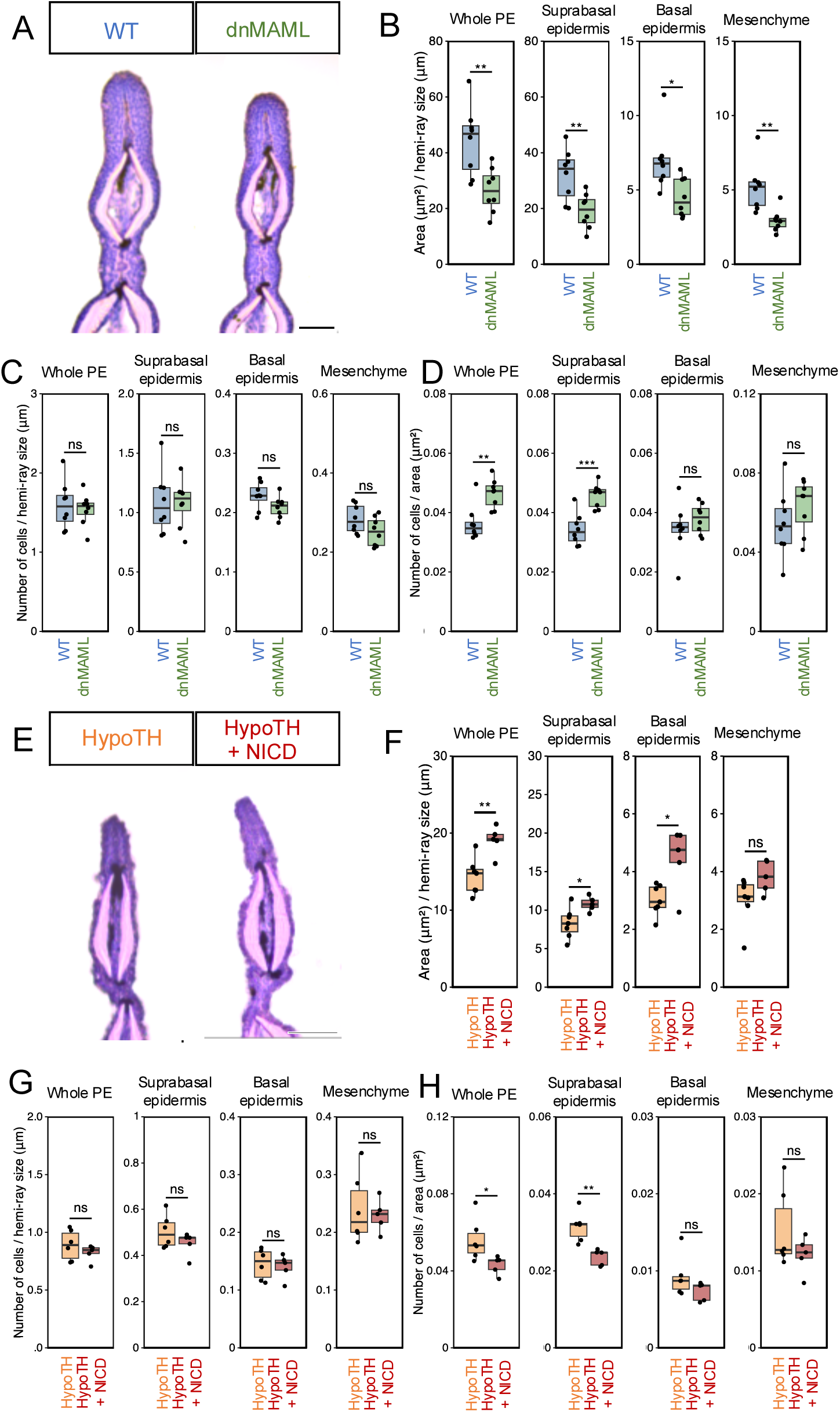
Notch pathway regulates PE growth. (A, E) Representative hematoxylin and eosin (HE) stained transverse sections of the peripheral regions. (A) compares wild-type (WT) controls with dnMAML-overexpressing fins, while (E) compares HypoTH controls with NICD-activated HypoTH fins. Scale bars: 0.5 mm. (B, F) Quantification of the normalized areas of whole PE and specific tissue layers within the PE, in dnMAML (B) and NICD (F) experiments. (C, G) Total cell number normalized to the hemi-ray size in dnMAML (C) and NICD (G) experiments. (D, H) Cell density (cell number per cross-sectional area) in dnMAML (D) and NICD (H) experiments. Data in (B-E, F-H) are presented as box plots showing the median, quartiles, and individual data points. Asterisks indicate significant differences between experimental groups (*P < 0.05, **P < 0.01; ns, not significant).

### Rescue with Notch activation partially rescues peripheral edge expansion in HypoTH backgrounds

To test if Notch activation is sufficient to rescue the reduced PE area of a HypoTH background, we induced overexpression of the Notch1a intracellular domain (NICD) in HypoTH fish. We crossed a heat-shock inducible NICD line^45^ into the HypoTH background. We confirmed the efficacy of this system by showing an upregulation of the Notch target *hey1* in the peripheral rays following heat shock (Fig. S7). Histological analysis revealed that sustained activation of the Notch pathway indeed increased the normalized PE area while preserving the stratified organization (Fig. 4E and F). Normalized areas of all tissue layers were significantly increased relative to those of HypoTH fish (Fig. 4F) and did not show significant changes in relative proportions (Fig. S8F and G). In NICD overexpressing HypoTH fins, cell number remained comparable to that in HypoTH controls (Fig. S8H and 4G), and a significant reduction in cell density was observed specifically in the suprabasal epidermis layer (Fig. S8H and 4H). The thickness of the IR region remained unaffected by Notch activation (Fig. S8I and J).

Taken together, these results suggest that Notch signaling is both necessary and sufficient to mediate the effects of TH in regulating expansion of the PE, supporting a model in which Notch signaling promotes PE growth primarily by inducing cell growth. By regulating cell expansion, Notch signaling ensures the proper growth and layer organization of the PE (Fig 5). We conclude that Notch serves as a critical downstream effector by which TH controls the hypertrophic growth of the PE.

**Figure 5.**
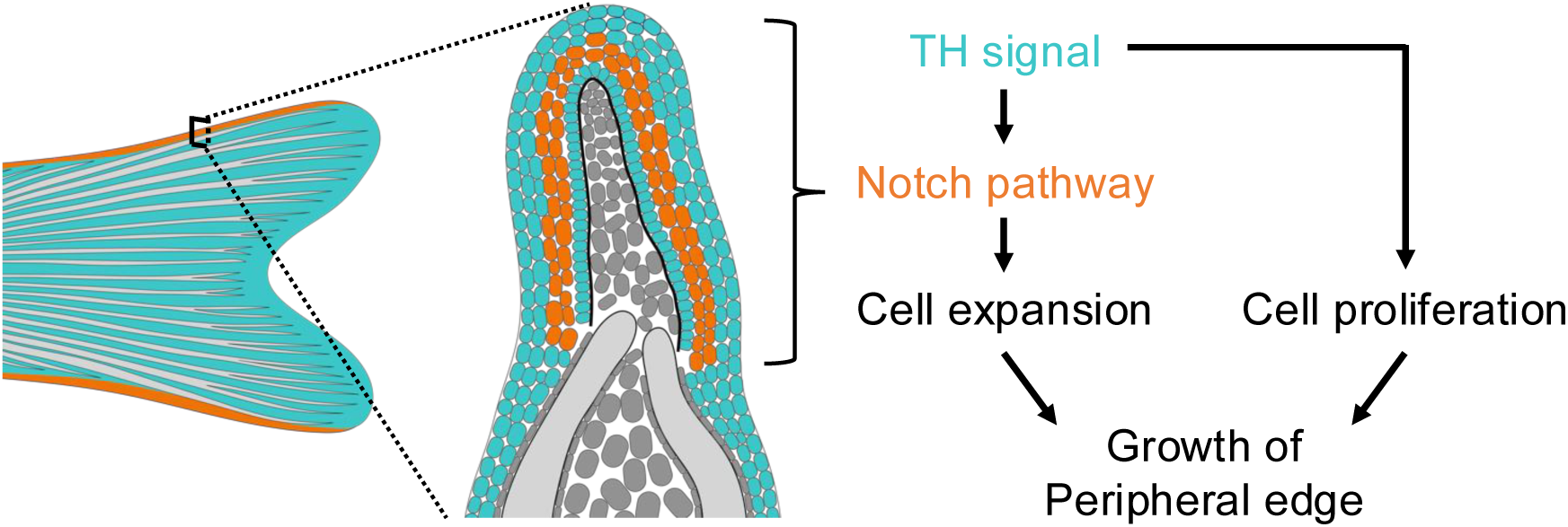
Proposed model for the regulation of PE growth by TH and Notch. Schematic showing the spatial domains of TH signaling (cyan) and Notch activity (orange). In this proposed model, TH signaling facilitates PE growth through cell proliferation and Notch-mediated cell expansion.

## Discussion

Using the zebrafish caudal fin as a model, we revealed a novel role for TH in vertebrate skin development. While TH regulates both cell proliferation and hypertrophy, these two processes are distinctly regulated; specifically, TH drives the hypertrophy of suprabasal cells via the Notch signaling pathway. Notably, although the molecular machinery for TH signaling is ubiquitously expressed throughout the fin epidermis, TH-responsive Notch activity is highly enriched in the PE and regulates its allometric expansion. Ultimately, these findings allow us to propose a model (Fig. 5) in which a systemic endocrine signal governs local structural morphogenesis.

Comparison of the PE and IR structures between juvenile and adult fins revealed that while the fundamental epidermal architecture is maintained, the PE and IR each adopt distinct growth strategies. Relative to fin ray width, the IR grows isometrically, whereas the PE exhibits positive allometric growth. Crucially, HypoTH fins failed to exhibit this positive allometric growth, suggesting that TH signaling is required to drive the normal expansion of the PE. We propose that this TH-dependent allometry is achieved through Notch signaling; our data showed that HypoTH fins exhibit reduced Notch activity. Consistent with this, continuous Notch inhibition reduced the PE area by decreasing cell size, partially phenocopying the HypoTH fins. Conversely, continuous Notch activation in HypoTH fins partially rescued the PE area by increasing cell size. Importantly, neither treatment affected IR thickness. These findings suggest that TH induces allometric growth of the PE, but not the IR, via the Notch pathway.

TH machinery was found to be broadly expressed throughout the fin epidermis. Despite this widespread competence to respond to systemic TH, Notch activity was predominantly enriched in the PE epidermis. We ruled out the idea that hydrodynamic stress experienced by the fin edge during swimming triggers this local Notch activation. Our findings support the model that the spatial patterning of Notch activity is governed by intrinsic tissue identity rather than extrinsic physical factors.

We propose that distinct positional information in the peripheral edge allows TH to locally activate Notch and PE expansion. Notably, previous proteomic data analyses revealed distinct protein expression profiles along the peripheral-central axis^46^. Consistent with this molecular heterogeneity, transgenic reporters for *lhx9* and *alx4a* are known to mark the peripheral rays in the pectoral fins^47,48^. It is highly likely that TH engages in crosstalk with these regional identity factors to induce the expression of Notch ligands, creating a permissive environment where Notch signaling is activated specifically in the PE. Systemic TH does not universally activate Notch; instead, it acts as a permissive signal interacting with local positional cues to drive region-specific growth.

In mammals, including humans, hypothyroidism is known to lead to severe cutaneous symptoms, including epidermal thinning and xerosis^10,49^. However, the molecular mechanisms underlying these pathologies—specifically how TH regulates skin homeostasis and structural maturation—have remained poorly understood. Here, we demonstrated that TH-deficient zebrafish exhibit distinct defects in epidermal maturation, leading to poor growth in the epidermis due to compromised Notch signaling.

Together, these findings suggest that a fundamentally conserved TH-Notch axis may play a broadly conserved role in regulating skin thickness and integrity. Indeed, targeted inhibition of Notch signaling in mice results in a significant reduction in epithelial thickness^50^. However, it remains to be determined whether TH acts upstream of Notch to regulate this process in mammals. Therefore, while the anatomical complexity of the skin varies across species, uncovering this hierarchical TH-Notch network provides a fundamental mechanistic framework. Future investigations into the potential conservation of this axis will be crucial for understanding the pathogenesis of endocrine-related skin atrophy in other vertebrates.

## Acknowledgements

We thank the entire McMenamin Lab for discussion and support, and in particular Tony Liu for his assistance with pilot experiments and initial data collection for the fin margin amputation assays. We also thank Melissa McTernan for statistical consultation and advice. This work was supported by the National Institutes of Health (NIH) under award number R35GM146467 (MIRA) to S.M.

## Methods

## Key resources table

| REAGENT or RESOURCE | SOURCE | IDENTIFIER |
| --- | --- | --- |
| **Antibodies** |  |  |
| Rabbit MCT10 (SLC16A10) Antibody | Thermo Fisher Scientific | Cat# PA5-48538, RRID:AB_2633995 |
| Rabbit DIO1 Antibody | Proteintech | Cat# 11790-1-AP, AB_10949197 |
| Rabbit DIO2 Antibody | Antibodies Online | Cat# ABIN652664 |
| Rabbit THRa Antibody | Genetex | Cat# GTX25621, RRID:AB_380602 |
| Mouse GFP Antibody | DSHB | Cat# N86/8, RRID:AB_2313651 |
| Mouse PH3 Antibody | Santa Cruz Biotechnology | Cat# sc-374669, AB_11150094 |
| Alexa Fluor 594 anti-rabbit IgG (H+L) | Molecular Probes | Cat# A11012, RRID:AB_141359 |
| Alexa Fluor 594 anti-mouse IgG (H+L) | Thermo Fisher Scientific | Cat# A11032, AB_2534091 |
| **Chemicals, peptides, and recombinant proteins** |  |  |
| TRIzol™ Reagent | Thermo Fisher Scientific | Cat# 15596026 |
| SuperScript IV First-Strand Synthesis System | Thermo Fisher Scientific | Cat# 18091300 |
| PowerUp SYBR Green Master Mix | Thermo Fisher Scientific | Cat# A25741 |
| **Critical commercial assays** |  |  |
| Direct-zol RNA Microprep Kit | Zymo Research | Cat# R2060 |
| **Experimental models: Organisms/strains** |  |  |
| *Tg(tg:nVenus-v2a-nfnB)* | McMenamin et al., 2014<sup>5</sup> | ZDB-ALT-141218-1 |
| *Tg(Tp1bglob:eGFP)* | Parsons et al., 2009<sup>44</sup> | ZDB-ALT-090625-1 |
| *Tg(hsp70l:DN-MAML-GFP)* | Zhao et al., 2014<sup>45</sup> | ZDB-ALT-140522-4 |
| *Tg(hsp70l:1xMYC-notch1a-intra)* | Zhao et al., 2014<sup>45</sup> | ZDB-ALT-140522-5 |
| **Oligonucleotides** |  |  |
| PCR primer sequences | IDT | See Table S1 |
| HCR split-initiator probes (B1 initiators) | Molecular Instruments | See Table S1 |
| B1 DNA HCR amplifiers | Molecular Instruments | N/A |
| **Software and algorithms** |  |  |
| Fiji | Schindelin et al., 2012<sup>51</sup> | RRID:SCR_002285 |
| Olympus cellSens Dimension | Olympus | RRID:SCR_014551 |
| Leica Application Suite X | Leica | RRID:SCR_013673 |
| R Project for Statistical Computing | The R Foundation | V4.4.1<br>RRID:SCR_001905 |
| RStudio | Posit | v2024.04.2+764<br>RRID:SCR_000432 |
| Diomni Design and Analysis software v3.0.1 | Thermo Fisher Scientific | v3.0.1 |
| fastp | Chen et al., 2018<sup>52</sup> | RRID:SCR_016962 |
| STAR | Dobin et al., 2013<sup>53</sup> | v2.7.10b<br>RRID:SCR_004463 |
| RSEM | Li et al., 2011<sup>54</sup> | v1.3.3<br>RRID:SCR_000262 |
| DESeq2 | Love et al., 2014<sup>55</sup> | V1.44.0<br>RRID:SCR_015687 |
| clusterProfiler | Xu et al., 2024<sup>56</sup> | 4.12.6<br>RRID:SCR_016884 |
| rstatix | Kassambara, 2023 | V0.7.2<br>RRID:SCR_021240 |
| Companion to Applied Regression | Fox and Weisberg 2019<sup>57</sup> | v3.1.3<br>RRID:SCR_022137 |
| Easy_HCR | Elagoz et al., 2022<sup>58</sup> | https://github.com/SeuntjensLab/Easy_HCR |
| org.Dr.eg.db | Carlson 2024<sup>59</sup> | https://bioconductor.org/packages/org.Dr.eg.db |

## Experimental model and subject details

### Fish lines and rearing conditions

All experiments with zebrafish were done in accordance with protocols approved by the Boston College Institutional Animal Care and Use Committee (protocol number 2007-006-01). HypoTH and their WT controls were both *Tg(tg:nVenus-v2a-nfnB)*^5^. The other transgenic strains used were: *Tg(Tp1bglob:eGFP)*^44^*, Tg(hsp70l:dnMAML-GFP)*^45^, *Tg(hsp70l:1xMYC-notch1a-intra)*^45^.

All fish were maintained at 28°C under a 14:10 light:dark cycle and were fed 2-3 times per day. Animals were considered juveniles when they were 11–13 mm standard length (SL) and considered to be adults when they were 18–25 mm SL. All larvae were fed live marine rotifers and *Artemia*. Juvenile and adult fish stocks that were not thyroid-hormone-modulated were fed Gemma Micro (Skretting) and Adult Zebrafish Diet (Zeigler). To minimize the potential introduction of exogenous TH, HypoTH adults and their WT controls were fed pure Spirulina flakes (Pentair) and live Artemia instead of the enriched pellet diet.

## Method details

### Thyroid Follicle Ablations

Thyroid hormone-deficient (HypoTH) fish were generated via thyroid follicle ablation as previously described^5^. Briefly, fish were treated with 10 mM metronidazole (MTZ, Alfa Aesar) in 1% DMSO (Sigma-Aldrich), or 1% DMSO alone (as a control), overnight at 28°C, and subsequently washed with system water.

### Fin amputation

After anesthetizing fish with MS-222 (Syndel), the dorsal-most peripheral ray or central rays were amputated at the position of the distal tip of the adjacent procurrent ray. The lateral ray was excised using a #11 scalpel blade, and then regeneration was prevented by trimming the regenerated tissues every other day.

### Pharmacological treatments

Stocks of T4 (L-thyroxine; Sigma-Aldrich) were prepared in NaOH, and subsequently diluted into the fish system water to achieve a final concentration of 10 nM T4 with 388 nM NaOH. Vehicle controls were prepared by adding the corresponding amount of NaOH to system water. Water changes were performed at least every other day throughout the treatment period.

### Heat Shock Treatment

Heat shock treatments were performed using two different regimens depending on the experimental purpose. For chronic treatments during development, heat shock was performed once daily from 10 dpf to adulthood. For acute Notch modulation prior to qPCR analysis, adult fish were subjected to heat shock once daily for two consecutive days, and fin tissues were collected for RNA extraction 5 hours after the completion of the second heat shock. For both regimens, a recirculating water system equipped with a heater was utilized. Specifically, the water temperature was gradually increased from the baseline of 28°C to 37°C over 1.5 hours, maintained at 37°C for 1 hour, and subsequently cooled back to 28°C within 30 minutes.

### Cryosection

Caudal fins were collected from fish anesthetized with MS-222 and immediately fixed in 4% paraformaldehyde (PFA) in phosphate-buffered saline (PBS) overnight at 4°C. Following two 10-minute washes in PBS, fins were decalcified in 0.45 M ethylenediaminetetraacetic acid (EDTA) in PBS overnight at 4°C and washed twice more with PBS. For cryoprotection, samples were sequentially incubated in a graded sucrose/PBS series: 10% and 20% sucrose for 1 hour each, followed by 30% sucrose for more than 3 hours at 4°C. Fins were then incubated in a 1:1 mixture of 30% sucrose and optimal cutting temperature (OCT; Fisherbrand) compound overnight at 4°C. Samples were embedded in OCT and stored at -80°C. Sections (16 μm) were obtained using a Leica CM1950 cryostat, mounted onto Superfrost Plus microscope slides (Fisherbrand), dried at room temperature for at least 30 minutes, and stored at -20°C.

### Hematoxylin and Eosin Staining

Sections were initially washed in running tap water for 5 minutes. Samples were subsequently incubated in Mayer’s hematoxylin (Electron Microscopy Sciences) for 2 minutes, followed by a second wash in running tap water for 5 minutes. Sections were counterstained by incubation in eosin Y (Electron Microscopy Sciences) for 5 minutes. Dehydration and clearing of the sections were then performed sequentially: 90% ethanol for 5 seconds, 100% ethanol for 5 seconds (repeated three times). Finally, samples were mounted using glycerol.

### Immunohistochemistry staining

Sections were washed twice in PBS and once in PBST (PBS containing 0.1% Tween 20). Blocking solution (PBST containing 3% sheep serum; Sigma-Aldrich) was applied for 1 hour. Slides were then incubated with the following primary antibodies: anti-MCT10 (1:400), anti-DIO1 (1:400), anti-DIO2 (1:400), anti-THRa (1:400) in the blocking solution overnight at 4°C, whereas anti-GFP was incubated for 48 hours. After washing three times with PBS, sections were incubated for 1 hour at room temperature with secondary antibodies: donkey anti-rabbit Alexa Fluor 594 (1:800) or goat anti-mouse Alexa Fluor 594 (1:200). Finally, the slides were washed three times with PBS, counterstained with DAPI (Sigma-Aldrich), and mounted using Histodenz (Sigma-Aldrich).

### Hybridization chain reaction

Probes for Hybridization Chain Reaction (HCR) *in situ* hybridization were designed using the Easy_HCR-main utility. The designed HCR split-initiator probes (incorporating B1 initiators) and corresponding B1 DNA HCR amplifiers (hairpins) were purchased from Molecular Instruments. The hybridization, wash, and amplification buffers were prepared in-house according to the manufacturer’s specified formulations. Whole-mount HCR was performed on adult caudal fins with modifications to the manufacturer’s protocol. Briefly, fins were fixed in 4% PFA in PBS overnight at 4°C. Samples were washed with PBST, treated with 3% H_2_O_2_ in PBST for 30 minutes at room temperature (RT), dehydrated in 100% methanol, and permeabilized in a 1:1 mixture of methanol and acetone overnight at -20°C. After rehydration with PBSTx, tissues were treated with Solution 1.1^60^ (8–10% aminoalcohol *N*,*N*,*N*′,*N*′-Tetrakis (2-hydroxyethyl) ethylenediamine; Sigma-Aldrich, 5% Triton X-100, and 5% urea; Sigma-Aldrich) for 1 hour at 37°C. Following washes with 5xSSCTx and 5xSSCT (saline-sodium citrate buffer containing 0.1% Triton X-100 or 0.1% Tween 20, respectively), samples were pre-hybridized in probe hybridization buffer for at least 1 hour at 37°C, and then incubated with the probe solution overnight at 37°C. Post-hybridization washes consisted of four 15-minute washes in probe wash buffer at 37°C, followed by washes in 5xSSCT at RT. Samples were pre-amplified in amplification buffer for at least 1 hour at RT, then incubated in the amplification solution overnight at RT. Final washes were performed extensively in 5xSSCT at RT before imaging. The detailed sequences of the HCR probes designed for this study are listed in Table S2.

### Mitosis analysis

Mitosis analysis was performed using a protocol adapted from Stoick-Cooper et al., 2010^61^. Sections were washed three times in PBS and incubated in 2N HCl for 20 minutes at 37°C for antigen retrieval. After three immediate washes in PBS, sections were incubated in blocking solution (1% PBSTx containing 1% sheep serum) for at least 1 hour. Slides were then incubated with anti-PH3 primary antibody (1:200) overnight at room temperature. Following three 5-minute washes, three 30-minute washes, and three 60-minute washes with PBS, sections were incubated with goat anti-mouse IgG Alexa Fluor 594 secondary antibody (1:800) for 1 hour at room temperature. Slides were washed three times with PBS, counterstained with DAPI (Sigma-Aldrich), and mounted using Histodenz (Sigma-Aldrich). The number of PH3-positive cells was quantified from three sections per fin, and the average value was used to represent a single biological replicate.

### RNA-seq analysis

To evaluate the transcriptional profiles of TH and Notch signaling components, we re-analyzed raw fastq files from our previously published dataset^33^. Raw reads were pre-processed using fastp to remove adapter sequences and low-quality reads, and subsequently aligned to the zebrafish reference genome (GRCz11) using STAR. Transcript abundance and Transcripts Per Million (TPM) were quantified using RSEM. Ensembl gene IDs were mapped to gene symbols using the org.Dr.eg.db R package, and counts/TPM values for redundant symbols were aggregated by summation. For the analysis of Notch pathway components, we applied a stringent filtering criterion: genes were included only if they exhibited a TPM ≥ 1 across all three biological replicates (n=3) in either the WT or HypoTH group. Differential expression analysis was performed using DESeq2 on rounded aggregated counts. A gene was defined as significantly differentially expressed if it met the threshold of an adjusted *P*-value < 0.05. Gene Set Enrichment Analysis (GSEA) was conducted using the clusterProfiler package to assess the global shift of the Notch signaling pathway. A ranked gene list was generated by sorting all genes based on their log_2_ fold change values. Enrichment scores were calculated against the KEGG zebrafish dataset (dre), specifically targeting the Notch signaling pathway (dre04330). For visualization, relative expression was calculated by normalizing the TPM of each sample to the mean of the WT group.

### Quantitative reverse transcriptase PCR

For WT, HypoTH, and T4-treated HypoTH fish, fin tissues were isolated from both the peripheral region (one ray each from the dorsal-most and ventral-most margins) and the central region (four rays) of the caudal fin. These samples were homogenized in TRIzol Reagent, and total RNA was extracted using the Direct-zol RNA Microprep Kit. For heat-shocked fish (dnMAML, NICD, and their respective controls), fin tissues were isolated exclusively from the peripheral region, homogenized in TRIzol, and extracted using a standard TRIzol chloroform extraction protocol.

cDNA was synthesized using SuperScript IV Reverse Transcriptase. Quantitative PCR was performed using the PowerUp SYBR Green Master Mix according to the manufacturer’s instructions on QuantStudio 3 real-time PCR system (Thermo Fisher Scientific). Data were analyzed using Diomni Design and Analysis software v3.0.1. Gene expression levels were normalized to *beta-actin*. The primer sequences for b*eta-actin* were previously described^27^. The primer sequences used for *hey1* are listed in Table S1.

### Imaging

Live zebrafish were anesthetized with 0.02% MS-222 in fish water. Live imaging of whole fins, including the *Tg(Tp1bglob:eGFP)* as well as fins processed for Hybridization Chain Reaction (HCR) in situ hybridization, was conducted using a Leica M205 FCA stereomicroscope equipped with a Leica K8 camera and Leica Application Suite X software. For histological analysis, sections stained with hematoxylin and eosin were visualized and captured using a Leica DM 1000 LED upright microscope with a Flexacam c5 camera. Tissue sections subjected to immunostaining or DAPI counterstaining were imaged using an Olympus IX83 inverted microscope equipped with a Hamamatsu ORCA Flash 4.0 camera. For these immunostained and DAPI images, extended focal imaging (EFI) and subsequent deconvolution were performed during acquisition using Olympus cellSens Dimension software. All acquired images were processed using Fiji; adjustments to contrast, brightness, and color balance were performed only within the linear range of the data to ensure that quantitative conclusions were not compromised by image adjustment.

### Image Quantification

All acquired images were quantified using Fiji^51^. Given that fin ray thickness varies along the proximal-distal axis^62^ and that fin size scales isometrically with overall body growth^63^, absolute measurements of the PE area could potentially vary depending on the sectioning position and the size of the individual fish. To account for these potential variables, we normalized the PE area to the cross-sectional area of the hemi-ray from the third-most peripheral fin ray (Fig. 1A).

For the quantification of Notch activity, all samples were imaged under identical acquisition settings. GFP fluorescence intensity of the *Tg(Tp1bglob:eGFP)* transgenic line was measured to evaluate reporter activity. To eliminate non-specific background signals, the maximum background fluorescence intensity was first determined for each image, and a signal threshold was applied to exclusively capture signals above this baseline. The region of interest (ROI) was anatomically defined as the PE region distal to the procurrent ray of the dorsal-most peripheral ray. Within this specific ROI, the total fluorescence intensity and the corresponding area were measured. Finally, the mean fluorescence intensity was calculated by dividing the total intensity by the ROI area.

### Statistical analysis

For all comparisons between two groups, Welch’s t-test was performed to accommodate potential unequal variances. For comparisons among four groups, Welch’s ANOVA was used, followed by post hoc pairwise Welch’s t-tests with Holm’s P-value adjustment.

For the qPCR data, a mixed-design ANOVA (Mixed ANOVA) was employed to account for both the within-subject factor (paired samples from peripheral and central regions obtained from the same individual fish) and the between-subject factor (genotype/treatment condition). Post hoc analyses for the mixed model involved targeted pairwise t-tests (paired t-tests for regional comparisons and independent t-tests for condition comparisons), with multiple testing corrections applied via the Holm method. All statistical analyses and graph generation were performed using R (version 4.4.1) in RStudio, utilizing the *rstatix* and *car* packages for statistical computing, alongside *ggplot2*, *dplyr*, *tidyr*, and *patchwork* for data visualization and analysis.

---

**Figure S1.**
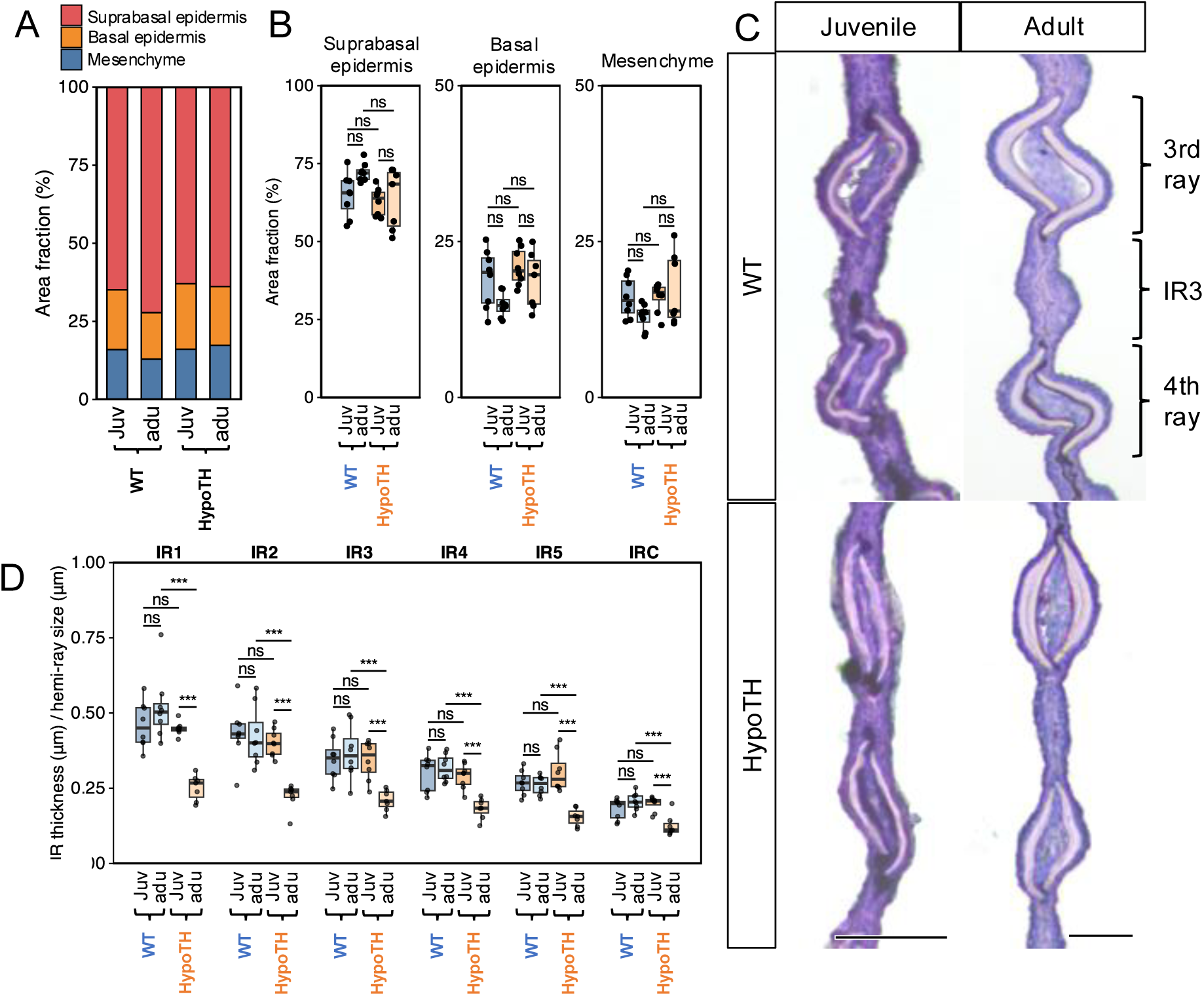
Tissue composition of the PE and thickness of the IR regions. **(A)** Area fractions of the suprabasal epidermis, basal epidermis, and mesenchyme within the total PE area. **(B)** Representative Hematoxylin and Eosin (HE) stained cross-sections of the third inter-ray region (IR3) in wild-type (WT) and TH-deficient (HypoTH) fish at juvenile and adult stages. Scale bars: 0.05 mm. **(C)** Quantification of the thickness of individual inter-ray regions (IR1–5 and IRC) normalized to the hemi-ray width. Data in (A) and (C) are presented as box plots showing the median, quartiles, and individual data points. Asterisks indicate significant differences (*P < 0.05, ***P < 0.001; ns, not significant).

**Figure S2.**
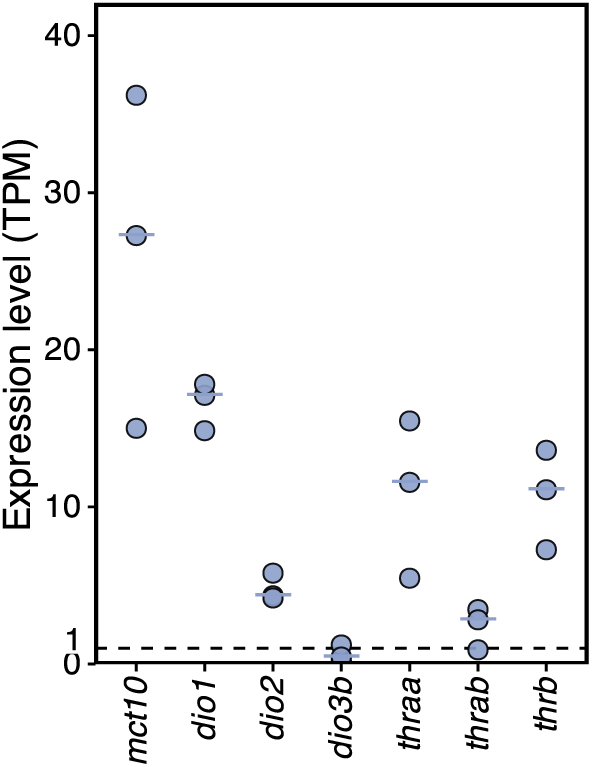
Gene expression levels of TH signaling pathway components. Expression levels of TH signaling pathway genes. Horizontal lines represent the mean. The dashed line represents TPM = 1.

**Figure S3.**
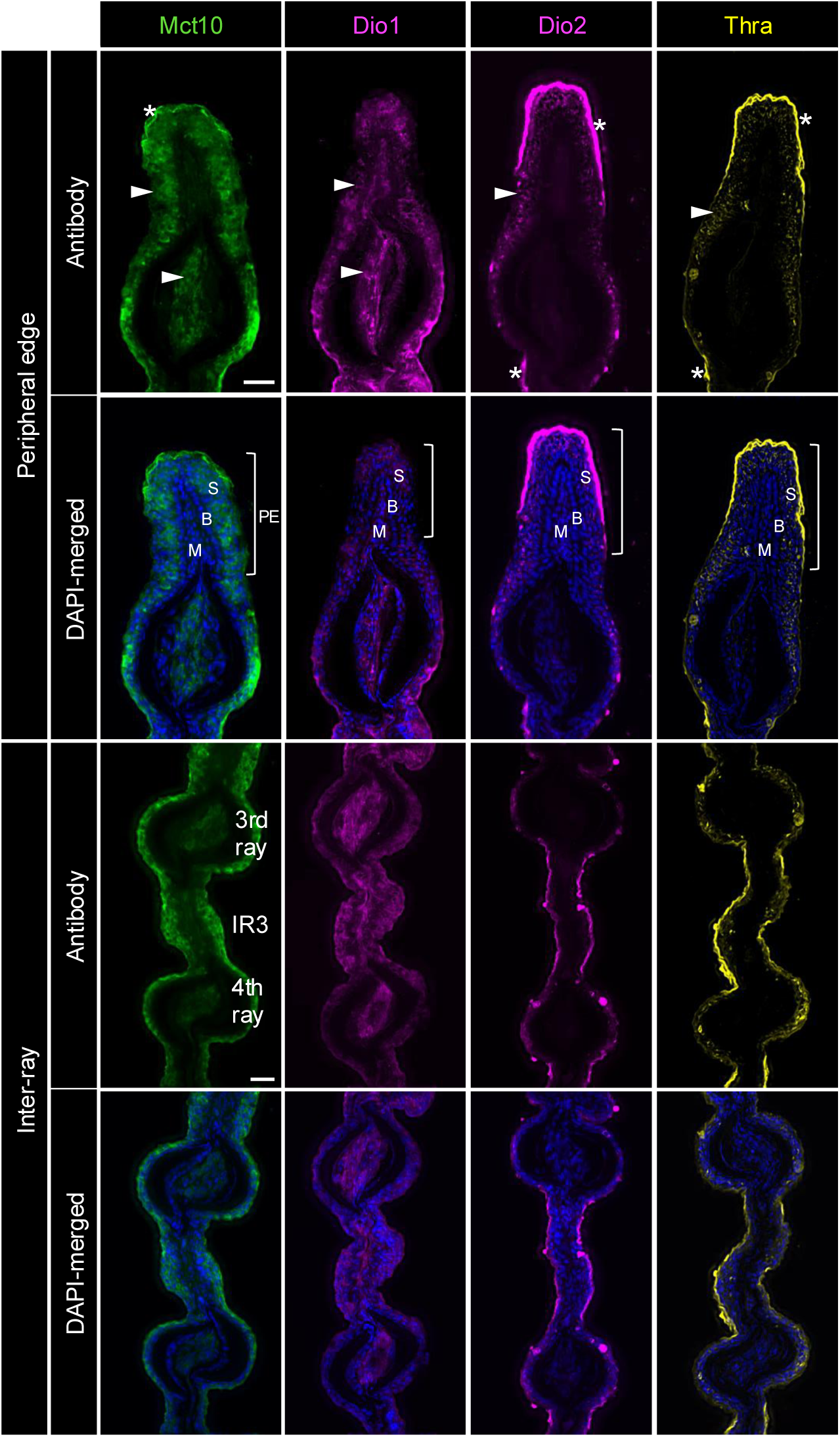
Localization of TH signaling components in the adult caudal fin. Immunohistochemical detection of Mct10, Dio1, Dio2, and Thra in cross-sections of the adult caudal fin. Panels display representative images of the peripheral edge (PE) and inter-ray (IR3) regions. Nuclei are counterstained with DAPI. White arrowheads indicate positive immunolocalizations. Asterisks mark background signals at the tissue margins. Scale bars: 20 µm. Abbreviations: S, suprabasal epidermis; B, basal epidermis; M, mesenchyme. (n = 3 per condition)

**Figure S4.**
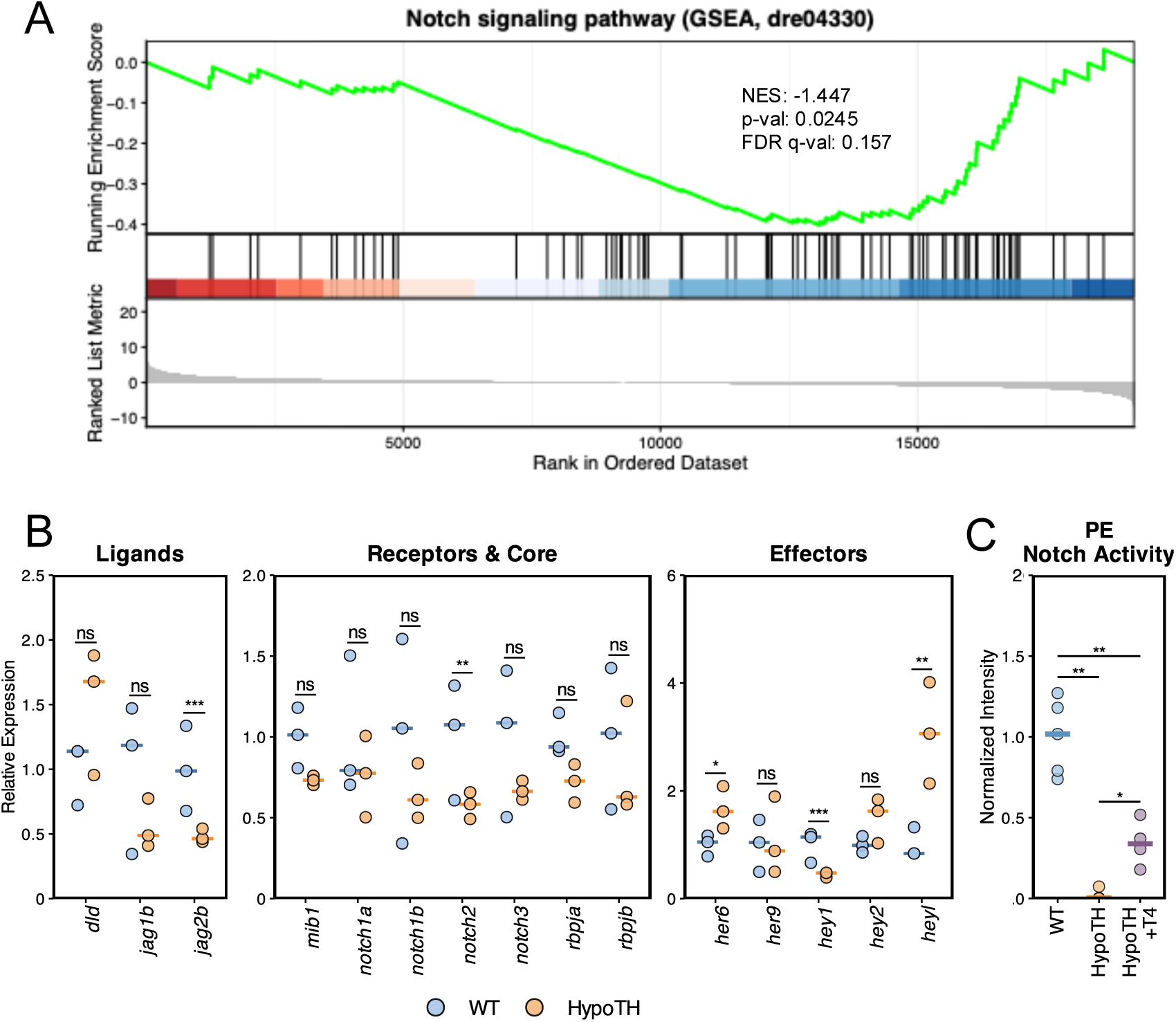
Thyroid hormone regulates Notch signaling activity and gene expression. (A) GSEA plot for the Notch signaling pathway. (B) Relative expression levels of Notch pathway genes in WT and HypoTH fin tissues. Horizontal lines represent the median. (C) Normalized GFP intensity in the PE (NaOH as vehicle control). Horizontal lines represent the median. Asterisks indicate significant differences (*P < 0.05, **P < 0.01, ***P < 0.001; ns, not significant).

**Figure S5.**
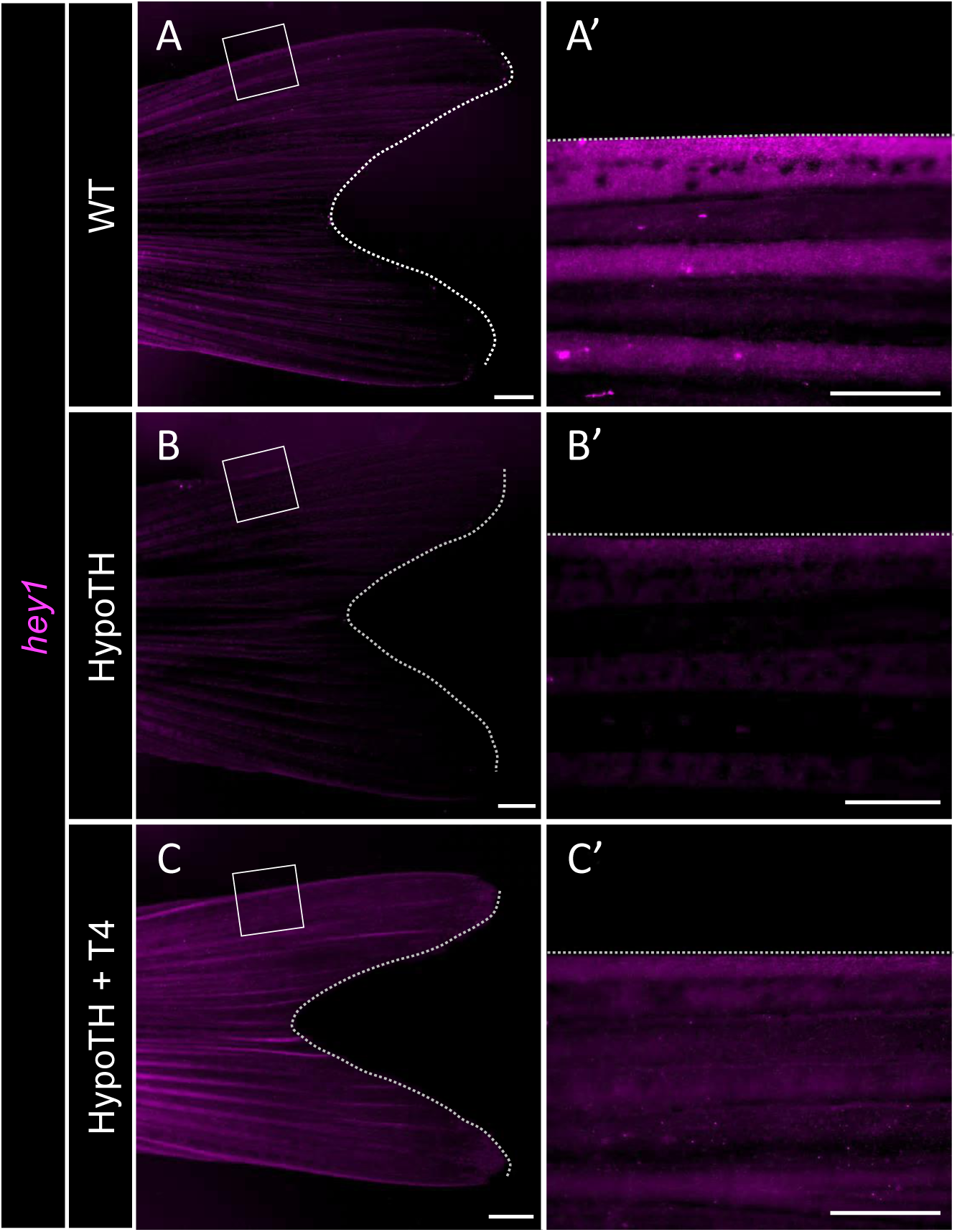
*hey1* expression pattern in the adult caudal fin. Representative images showing *hey1* expression detected by HCR in whole-mount fins (A–C) and corresponding magnified views (A’–C’) of the peripheral region for WT (A, A’), HypoTH (B, B’), and T4-treated HypoTH (C, C’) fins. Dotted lines mark the fin margins. Scale bars: (A-C) 1 mm, (A’-C’) 0.5mm. (n = 3 per condition)

**Figure S6.**
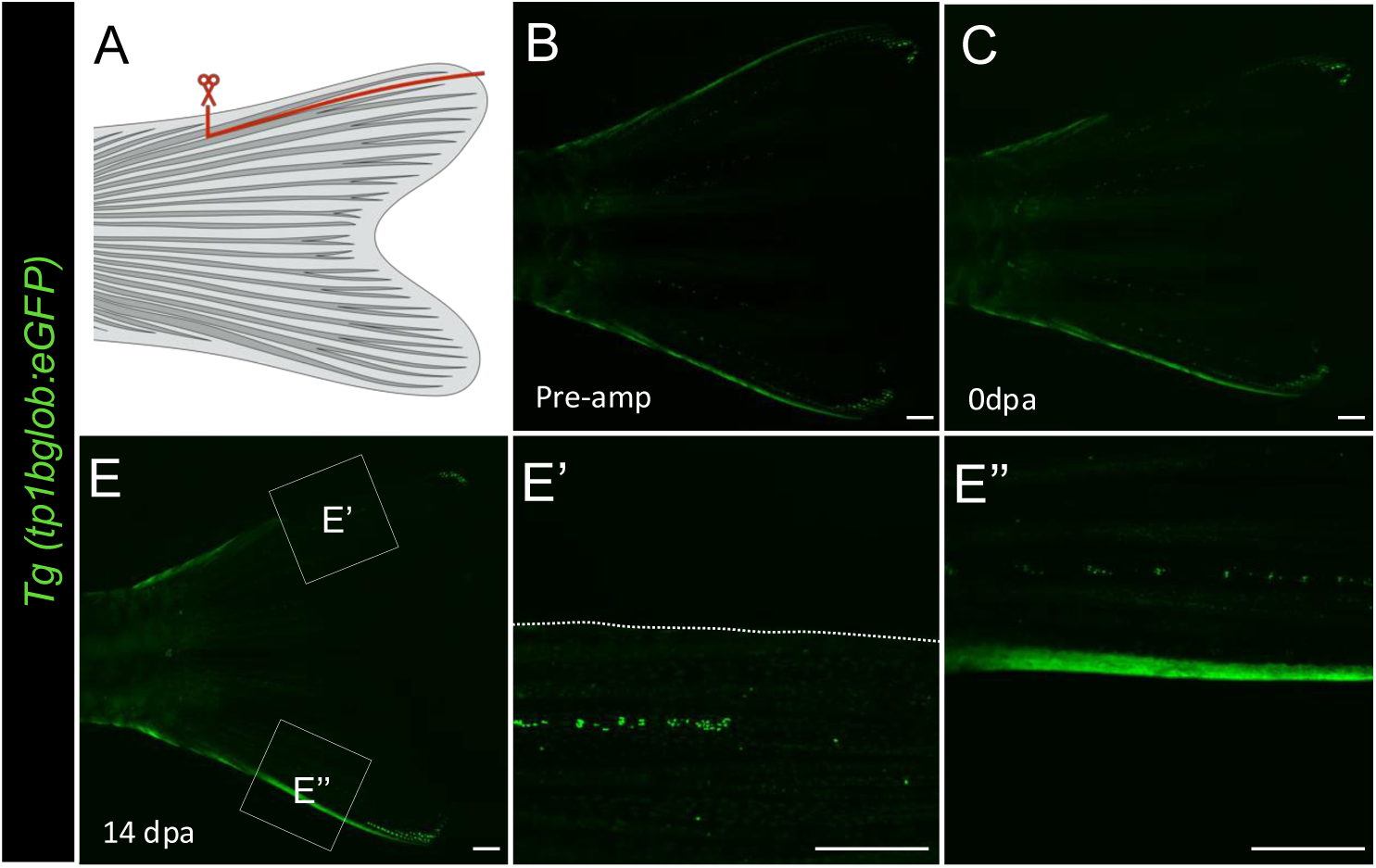
Assessment of Notch activity under increased mechanical stress. (A) Schematic of the amputation experiment. The red line indicates the removal of the most-peripheral ray. (B–E’’) Live imaging of *Tg(Tp1bglob:eGFP)* fins before (Pre-amp, B) and after (0 dpa, C; 14 dpa, E) amputation. (E’, E”) Magnified views of the peripheral edge of the adjacent rays indicated in (E). The dotted line in (E’) marks the fin margin. Scale bars: 1 mm. (n = 5 fins)

**Figure S7.**
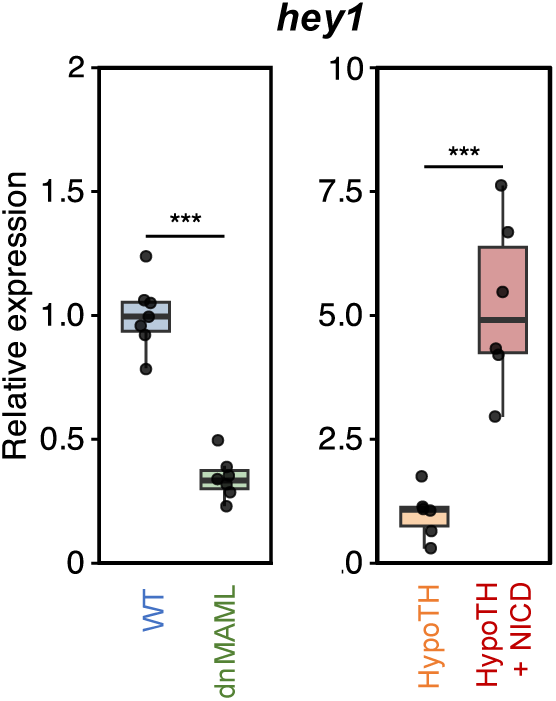
Transcriptional response of *hey1* to Notch pathway manipulation. qPCR analysis of *hey1* expression levels in peripheral fin tissues following heat-shock. Data are presented as box plots with individual data points. Asterisks indicate significant differences (***P < 0.001).

**Figure S8.**
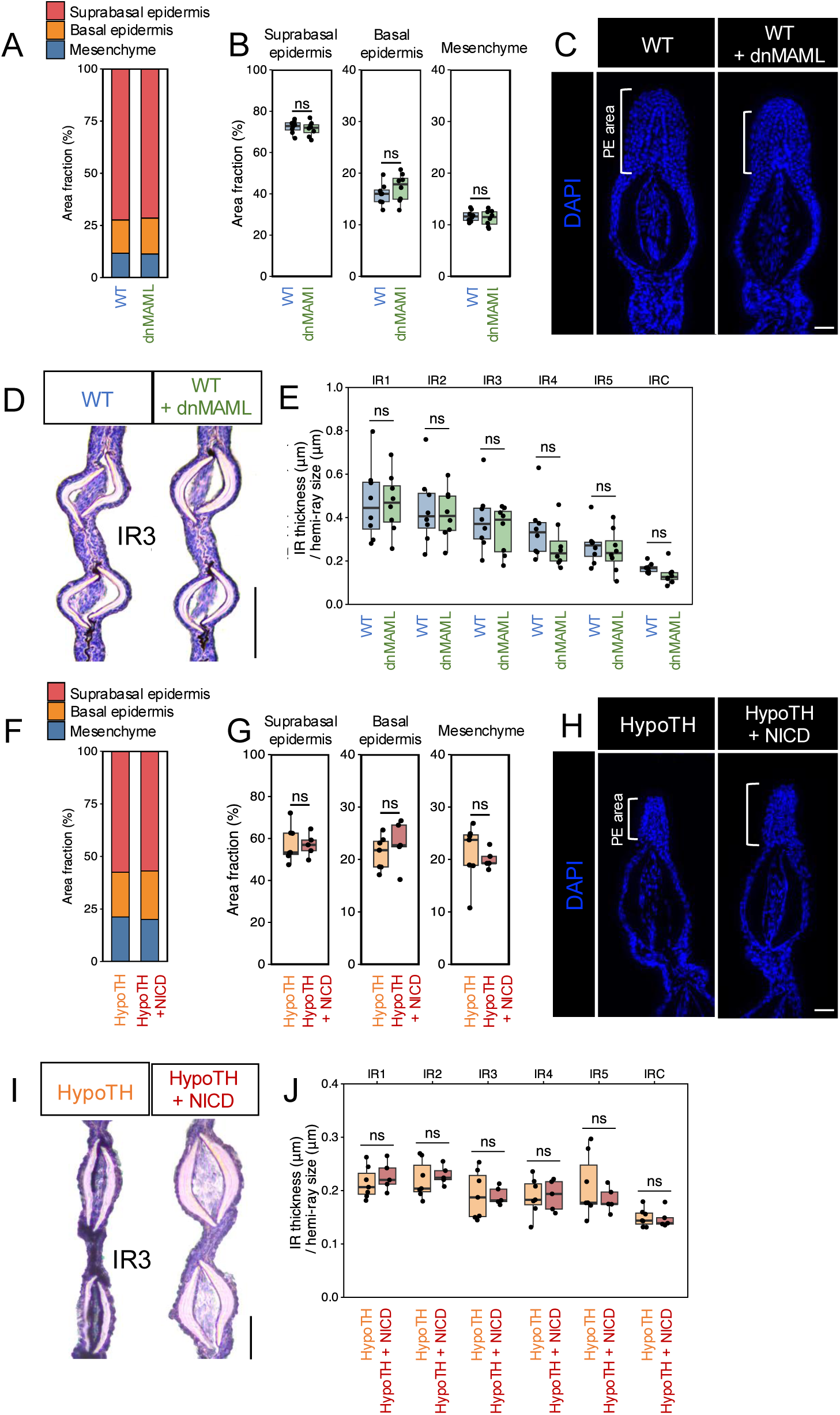
Tissue composition of the PE and thickness of the IR regions. (A, E) Area fractions of the suprabasal epidermis, basal epidermis, and mesenchyme within the total peripheral edge (PE) area in dnMAML (A) and NICD (E) experiments. (B, F) DAPI-stained cross-sections of the peripheral edge (PE). (B) compares wild-type (WT) controls with WT + dnMAML fins, while (F) compares HypoTH controls with HypoTH + NICD fins. Scale bars: 20 μm. (C, G) Representative hematoxylin and eosin (HE) stained cross-sections of the third inter-ray region (IR3) in dnMAML (C) and NICD (G) experiments. Scale bars: 0.5 mm. (D, H) Quantification of the thickness of individual inter-ray regions (IR1–5 and IRC) normalized to the hemi-ray width in dnMAML (D) and NICD (H) experiments. Data in (A, D, E, H) are presented as box plots showing the median, quartiles, and individual data points. ns, not significant.

**Table S1.**
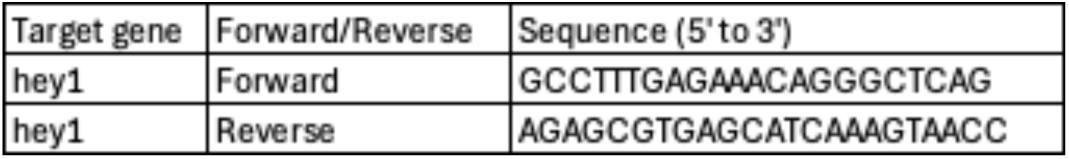
Primers used for qPCR.

**Table S2.**
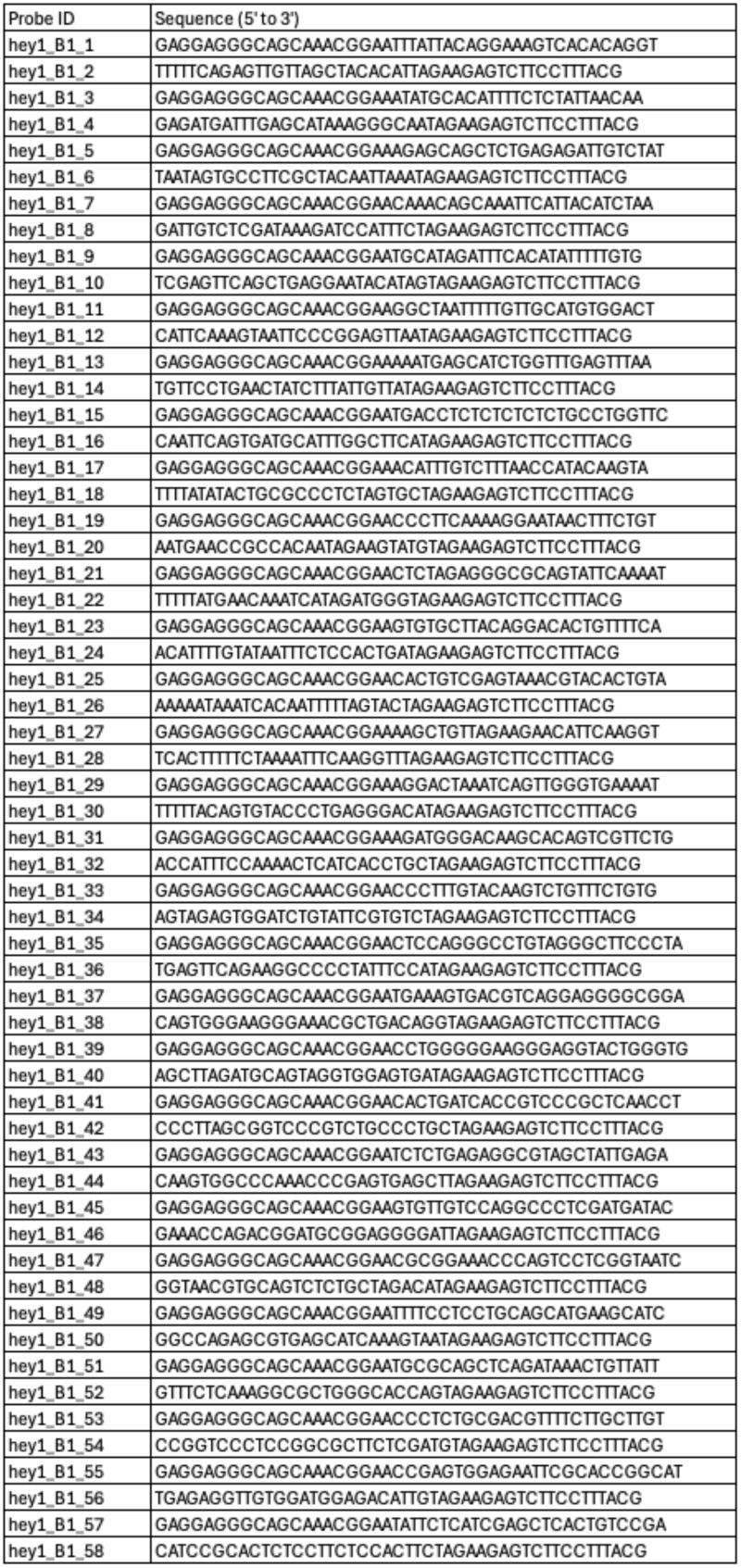
HCR probe sequences for *hey1*.

